# CellSNAP: A fast, accurate algorithm for 3D cell segmentation in quantitative phase imaging

**DOI:** 10.1101/2023.07.24.550376

**Authors:** Piyush Raj, Santosh Paidi, Lauren Conway, Arnab Chatterjee, Ishan Barman

**Author notes:** Ishan Barman –.

## Abstract

Quantitative phase imaging (QPI) has rapidly emerged as a complementary tool to fluorescence imaging, as it provides an objective measure of cell morphology and dynamics, free of variability due to contrast agents. In particular, three-dimensional (3D) tomographic imaging of live cells has opened up new directions of investigation by providing systematic and correlative analysis of various cellular parameters without limitations of photobleaching and phototoxicity. While current QPI systems allow the rapid acquisition of tomographic images, the pipeline to analyze these raw 3D tomograms is not well-developed. This work focuses on a critical, yet often underappreciated, step of the analysis pipeline, that of 3D cell segmentation from the acquired tomograms. The current method employed for such tasks is the Otsu-based 3D watershed algorithm, which works well for isolated cells; however, it is very challenging to draw boundaries when the cells are clumped. This process is also memory intensive since the processing requires computation on a 3D stack of images. We report the CellSNAP (Cell Segmentation via Novel Algorithm for Phase Imaging) algorithm for the segmentation of QPI images, which outstrips the current gold standard in terms of speed, robustness, and implementation, achieving cell segmentation under 2 seconds per cell on a single-core processor. The implementation of CellSNAP can easily be parallelized on a multi-core system for further speed improvements. For the cases where segmentation is possible with the existing standard method, our algorithm displays an average difference of 5% for dry mass and 8% for volume measurements. We also show that CellSNAP can handle challenging image datasets where cells are clumped and marred by interferogram drifts, which pose major difficulties for all QPI-focused segmentation tools. We envision our work will lead to the broader adoption of QPI imaging for high-throughput analysis, which has, in part, been stymied by a lack of suitable image segmentation tools.

## Introduction

Regulation and coordination of cell shapes are central to native physiology during all stages of organismal existence. Recent advances in optical imaging have provided mechanistic insights into such phenomena by revealing cellular features and processes with previously unimagined detail [1] [2] [3]. Central to the accurate analysis of such intricate biological processes is the precise segmentation of cellular images. Quantifying cellular morphology, like shape, area, circularity, aspect ratio, and others, starts with first segmenting the cells in a given field of view. Due to its unquestionable significance, much work has been done to standardize the process. There are well-developed open-source software suites, notably CellProfiler [4] and CellPose [5], to perform such segmentation tasks with great accuracy. The recent update to CellProfiler includes the functionality of 3D image segmentation and is currently the most widely used tool to perform such tasks. However, since the current workhorse for biological imaging is fluorescence microscopy, all the standard segmentation software are optimized for and targeted to the analysis of fluorescence images.

Yet, contrast-agent-free microscopy is highly desirable to study the dynamics and physiological activity of various structures in living cells. Quantitative phase imaging (QPI) measures optical field images using laser-based interferometry and has rapidly emerged as a viable imaging alternative because it offers an objective measure of morphology and dynamics in a label-free manner [1]. In addition to the amplitude images provided by conventional intensity-based microscopy techniques, QPI measures optical phase delay maps governed by the refractive index (RI) distribution of a sample. Since the endogenous RI distribution is strongly related to the structural and biochemical characteristics of the cell type, the acquired field images can be analyzed for systematic discovery of cell type-specific morphological and biophysical fingerprints encoded in the images.

Over the past two decades, QPI has provided important insights into diverse biological phenomena, ranging from membrane dynamics of red blood cells [6] to neuronal activity [7] and cell-nanoparticle [8] [9], and cell-drug interactions [10]. Recently, it has also been shown that QPI images can be mapped to fluorescence images using deep learning techniques, a concept coined as image-to-image translation. Prediction of stains (i.e., where specific fluorophores/stains would bind in an unlabeled specimen) using a combination of QPI and machine learning has been successfully demonstrated [11] [12] [13], and gradually, more stains are being added to the library. Indeed, phase imaging with computational specificity has allowed precise measurements of the growth of nuclei and cytoplasm independently, over many days, without loss of viability.

Central to many of the aforementioned and other emergent applications is QPI’s intrinsic ability to measure single-cell volume and mass non-destructively and ultra-sensitively over arbitrary periods of time in both adherent and flowing cell populations [1]. A critical step in undertaking such analysis is the accurate segmentation of the tomographic images of the cell populations. Since QPI imaging is still a relatively new technique in the field of cell biology, the analysis pipeline is not as developed as it is for fluorescence images. The toolbox developed for fluorescence image segmentation does not work well with QPI images, as the fluorescence contrasts are much sharper than the refractive index contrast. Also, in some segmentation procedures, the stained nucleus is used as a fiducial marker to define the respective cytoplasm boundary, and consequently, such algorithms cannot be directly implemented in QPI imaging. This has motivated researchers to develop segmentation algorithms tailored for QPI images [14], yet, their applicability has been limited to two-dimensional images thus far.

The state-of-art method used for 3D QPI cell segmentation is an Otsu-based 3D watershed algorithm [15] (hereafter referred to as the Otsu threshold algorithm in this work). This algorithm works very well for isolated cell images; however, it is challenging to draw boundaries when the cells are clumped. This process is also memory intensive since the processing requires computation on a 3D stack of images. As a point of reference, using the current state-of-the-art software module, this cell segmentation takes *ca*. 10 seconds for a 3D stack of images [484 × 484 × 208] on a workstation equipped with an 8^th^ generation i7 processor running at 3.7 GHz, 64GB RAM, and Nvidia GeForce GTX 1080 - 8GB graphics card. Another recently developed method for 3D QPI cell segmentation is an AI-based cell segmentation tool [16]. While this should, in principle, be able to overcome the limitations of the Otsu threshold algorithm, such implementation would require an extensive training dataset with enough complexity to handle all the edge cases. The preparation of such training datasets would also require manual annotations, besides needing separate datasets for different cell types. While an AI-trained system can be implemented on a modest system with a GPU, the training for an AI-based segmentation tool would necessitate a state-of-the-art computational resource.

To overcome these drawbacks, we report here a fast and light-weight algorithm for 3D QPI image-based cell segmentation. Our cell segmentation algorithm is inspired by the process of gemstone extraction, where a stonemason starts by making a coarse extrusion cut along a given axis and then gradually carves away near the surface of gemstone to reveal the final form. The algorithm first extrudes a 3D image from a 2D segmented mask, which can be seen as creating a rough shape of the cell structure. A 2D image is generated through maximum intensity projection of the 3D image, and a 2D cell segmentation algorithm is used to make a 2D mask. This allows us to identify the boundary region for segmentation in the x-y plane. Next, we take advantage of the continuity of cells and the absence of abrupt changes in consecutive z-stacks of the 3D image to perform the complete 3D segmentation. This step can be likened to the chiseling process in gemstone extraction, where the stone mason uses finer tools to carve away at the surface. The implication for developing such a method is vast as it can speed up high-throughput single-cell analysis and reduce the computational burden on the hardware. This also lowers the barrier to the adoption of QPI in biological sciences. The growing advancements in hardware development for high-throughput imaging in Quantitative Phase Imaging (QPI) [17] [18] also offer significant utility to this method.

## Design principle and workflow

**Step 1**: The first stage involves gathering and exporting all unprocessed 3D image stacks from the imaging software. Our code is tailored to operate with a tiff stack file, which contains matrix values represented by refractive index, saved in uint16 format. Nevertheless, the algorithm is compatible with any file format provided that we are utilizing a scalar multiplication of the refractive index value as the matrix value.

**Step 2**: We produce a Maximum Intensity Projection (MIP) image of the 3D tiff image stack along the z-axis. MIP is a technique used to visualize 3D data along a visualization axis, where only the voxels with maximum intensity are projected onto the image. Originally developed for use in Nuclear Medicine by Jerold Wallis, it has since been applied to various tomographic imaging modalities, such as CT scans and X-ray imaging [19]. Most 3D imaging software offers the option to export 2D MIP images or a standard library can also be utilized for this image-processing step.

**Step 3**: In this step, we take advantage of already well-built 2D image analysis libraries, which work wonderfully on 2D images for cell segmentation. Specifically, we use CellProfiler [1] to perform 2D cell segmentation on MIP images. The image mask files were saved in a separate folder. While CellProfiler was our tool of choice because of its ease of usage and flexibility, any 2D segmentation tool can be used as long as they have the ability to generate mask images.

**Step 4**: The 2D mask generated from CellProfiler is then extruded to 3D, where the number of image stacks for the mask is equal to the number of the z-stack planes in the original refractive index image.

**Step 5**: The process begins by selecting the mask for each cell within the field of view individually and setting the matrix value for that mask to 1, while the background and the mask of all other cells are set to 0. The resulting matrix is then multiplied by the original refractive index image. This process is repeated iteratively for all cells in the given field of view. Additionally, all pixels with refractive index values equal to or lower than the background refractive index value are set to zero. The resulting matrix, referred to as the *rough_segmentation_matrix*, shows a rough outline of the cell with noisy pixels around it, as depicted in Figure 1.

**Figure 1:**
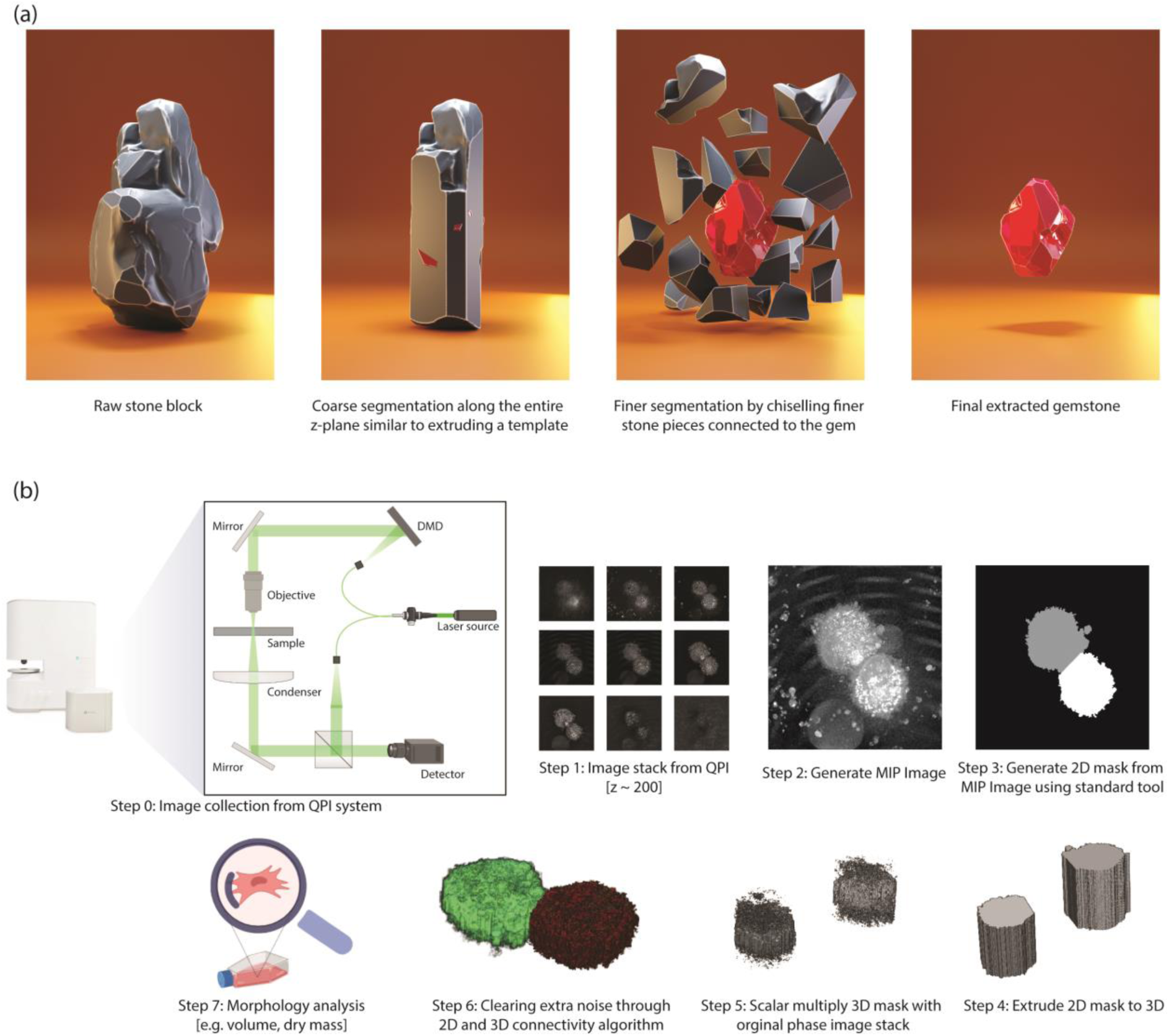
Our method draws inspiration from gemstone extraction from stone, where a stone block resembles a 3D raw image matrix. Coarse segmentation along the z-plane corresponds to step 4 in (b), while Fine segmentation aligns with step 5 and 6 in (b). (b) Step by step workflow for cell segmentation
using QPI images

**Step 6**: In this step, we use the continuity feature of 3D cell image and implement 2D and 3D connectivity matrices to eliminate noisy pixels around the cell. First, we take the *rough_segmentation_matrix* and plot the number of non-zero pixels (normalized) against the number of z slices. We set a hard threshold at the z-slice where the normalized non-zero-pixel value is 0.5. Instead of hard-coding the z-stack number to identify the bottom of the cell, we used a threshold condition because our microscope system has an axial resolution of 300 nm, which is lower than the surface roughness of commonly used petri dishes. Next, for each 2D plane, we label the connected components using the 2D connectivity matrix algorithm, achieved using the bwlabel function in MATLAB. We only retain the label corresponding to the highest number of pixels in each 2D slice plane to remove extra noise from cell debris (Fig. 2a). After 2D noise removal we use the bwlabeln function on the 3D image to eliminate all the noise in the 3D structure, removing all disconnected structures in the process, resulting in perfectly segmented cells (Fig. 2b).

**Figure 2:**
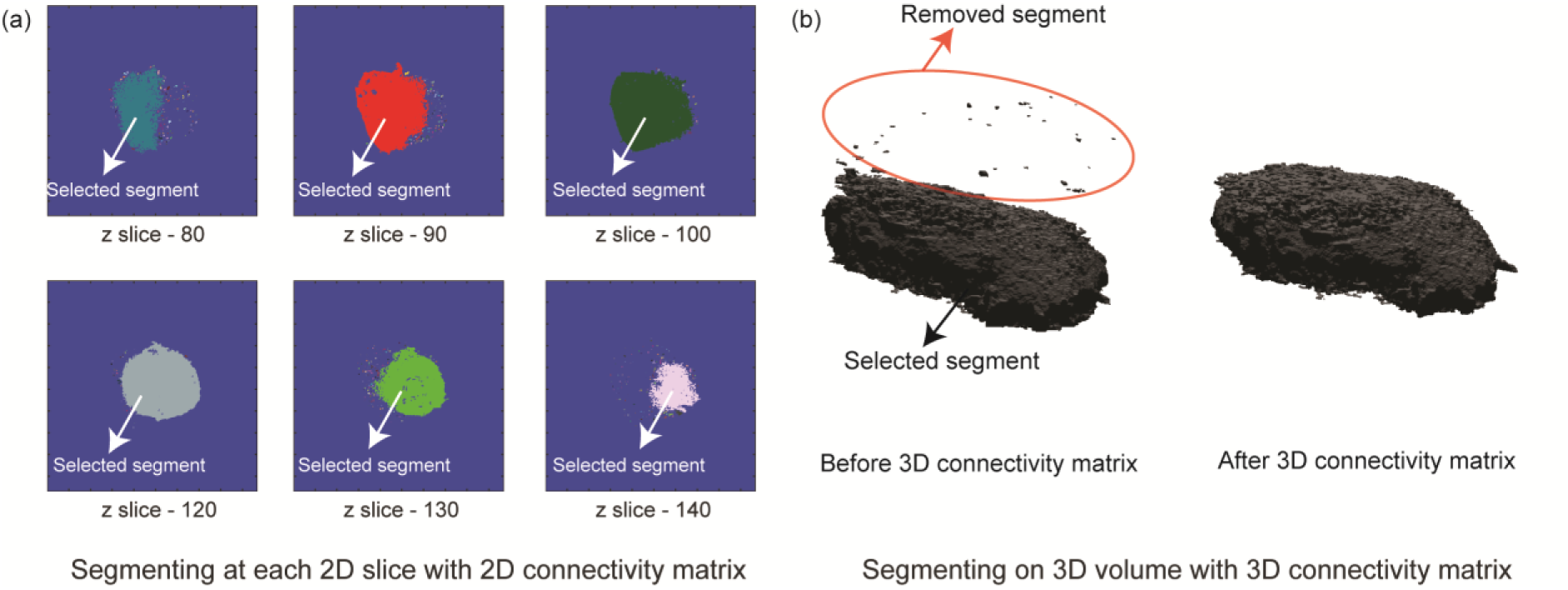
(a) The segmentation process involves utilizing a 2D connectivity matrix to label segments in each 2D slice, followed by the selection of the segment with the highest pixel count. Small segments, depicted in different colors withtin each slice, are subsequently discarded. (b) Analogously, the 3D segmentation process involves labelling distinct segments using a 3D connectivity matrix and selecting the segment with the highest voxel count.

**Step 7**: The dry mass is calculated using the following formula:

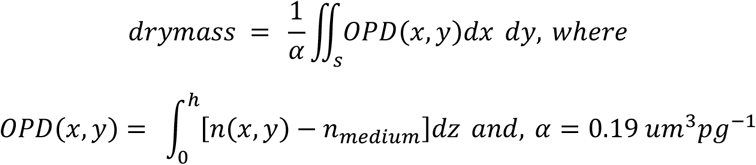

Volume is calculated by counting the number of non-zero pixels in the 3D segmentation mask.

## Results and Discussion

We tested our QPI cell segmentation algorithm against the current gold standard of the Otsu-based 3D watershed algorithm provided by the manufacturer with their microscopy software suite. We also used the recent AI-powered segmentation tool [20] for cases where the watershed algorithm did not provide satisfactory segmentation. The AI-powered segmentation tool is currently available for a limited number of cell types, including macrophages. Our comparative study, therefore, is with U937-derived macrophages M1 and M2 cells. Since label-free volume and dry mass measurements are unique to QPI images, these two metrics act as suitable benchmarking metrics to compare the performance of our algorithm. Cell experiments were conducted in sparsely populated cell populations in a petri dish, with a confluency of approximately 65%. This enabled us to obtain high-quality images featuring a single cell within a given field of view. As a result, we could compare our algorithm to the current gold standard with minimal concern for its accuracy, as the latter is plagued by robustness issues in the presence of multiple cells within a given field of view. We observed a dry mass difference of approximately 5% for both M1 and M2 cells, while the difference in volume was slightly higher, with a mean value of 8% for M1 and 6% for M2 cells. Since the methods used to perform cell segmentation differed significantly, it is encouraging to note that the mean differences in dry mass and volume were both less than 8%. Additionally, we have included a table that compares the computation time required by each of these algorithms, demonstrating that our algorithm significantly outperforms the current gold standard even on a platform with modest computational hardware.

We embarked on developing a cell segmentation algorithm due to the limitations of the current method in segmenting cells when they are clumped together within a field of view. Moreover, the current method’s performance is hampered in suboptimal imaging conditions. Given that QPI imaging is widely used for longitudinal studies, temperature drifts, stage drifts, and calibration drifts are inevitable, rendering the current segmentation algorithms unsuitable for such scenarios, with resulting imaging data being either unusable or requiring extensive manual processing. In this regard, we present two instances where our algorithm’s robustness is evident. In cases where multiple cells are clumped together and share boundaries, the existing Otsu thresholding fails to distinguish between the cells, resulting in a single unit being segmented (Fig. 4a). Although the AI segmentation tool offers some promise, as evidenced by the accurate segmentation of one cell in Fig. 4a, it fails to accurately segment the remaining cells. By contrast, our segmentation algorithm accurately segments the cells and outperforms existing methods in terms of computational speed.

**Figure 3:**
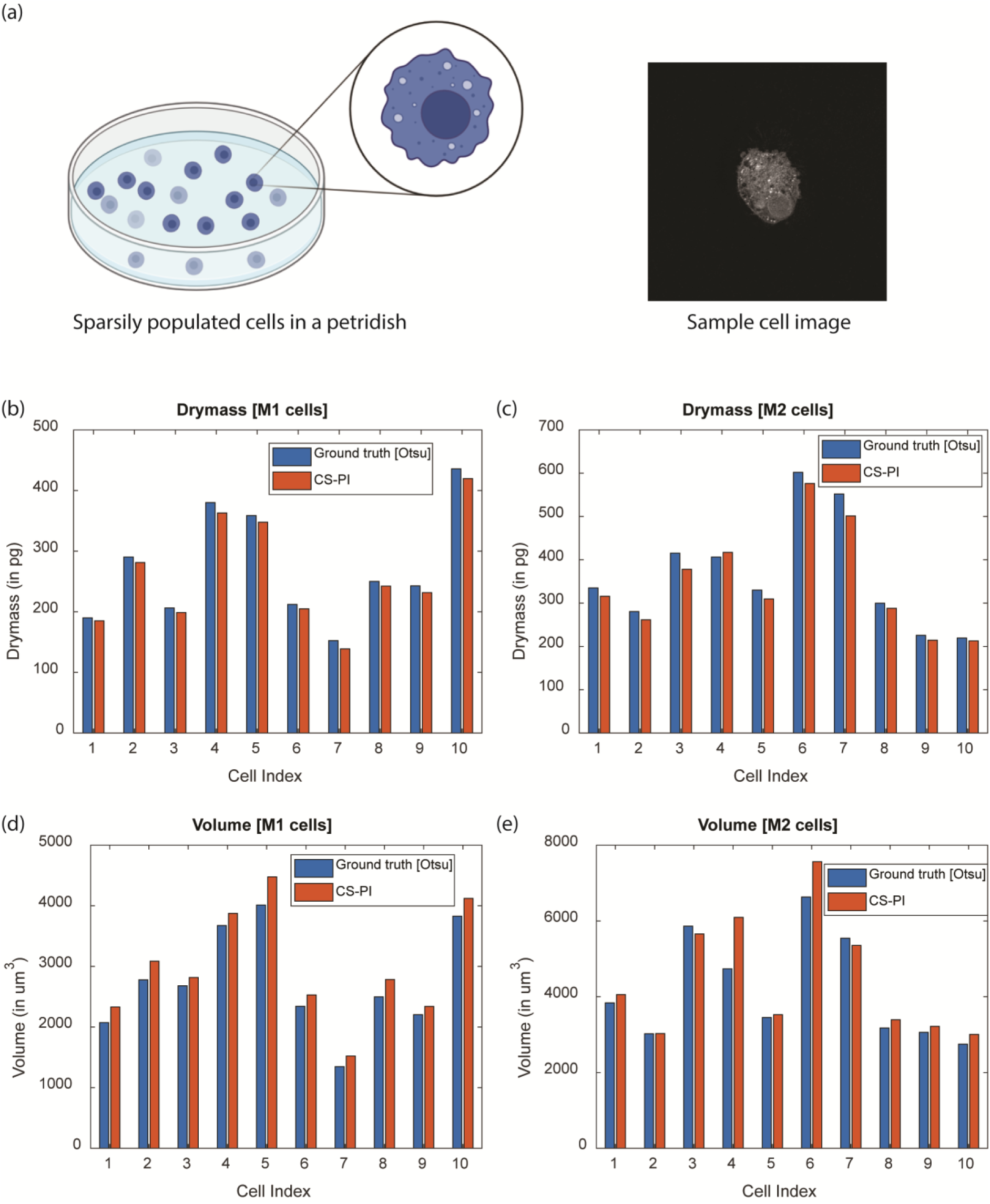
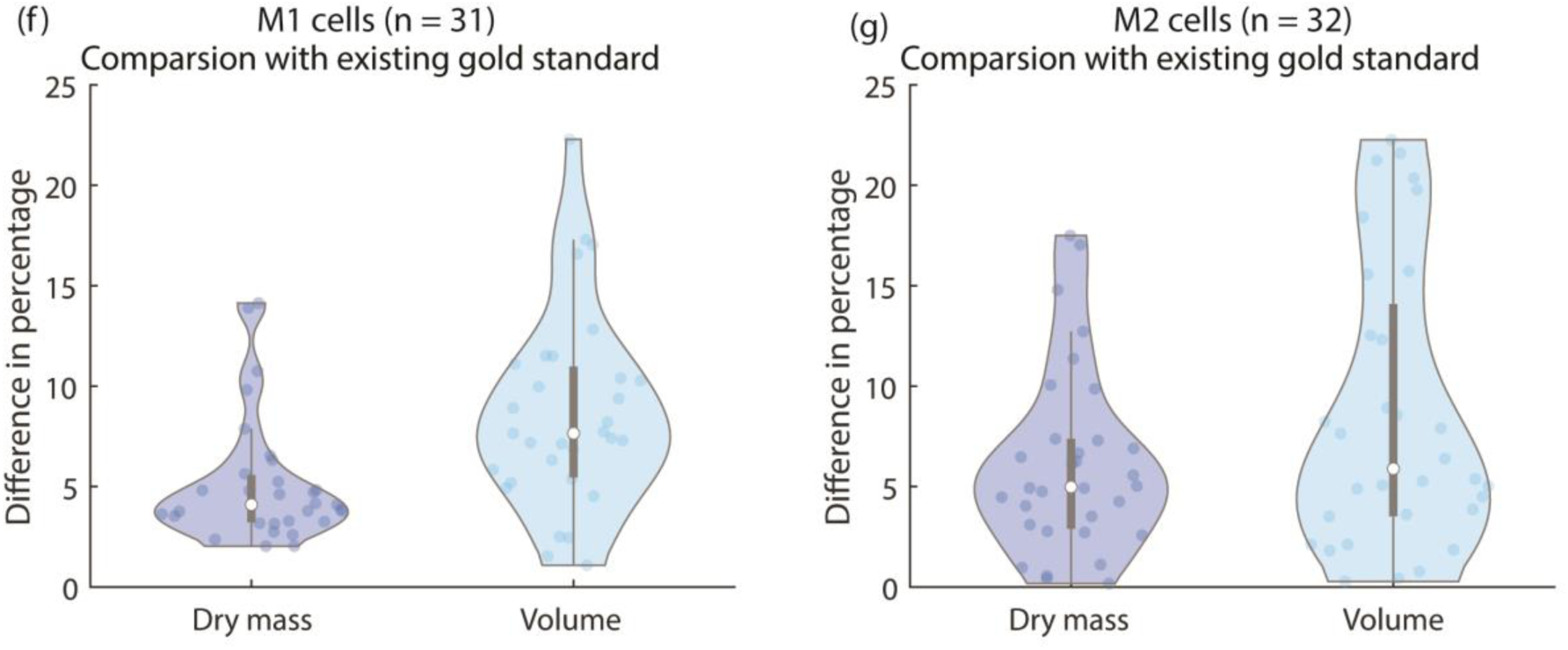
(a) Cell experiments with sparsely populated cells in the petridish and a sample cell image obtained using the QPI microscope. (b) Comparison of dry mass values for M1 cells. (c) Comparison of dry mass values for M2 cells. (d) comparison of volume for M1 cells. (e) Comparison of volume for M2 cells. For a statistical comparison between the two algorithms, the difference in the percentage of dry mass and volume was calculated for M1 (f) and M2 (g) cells with the current gold standard and our proposed algorithm.

**Figure 4:**
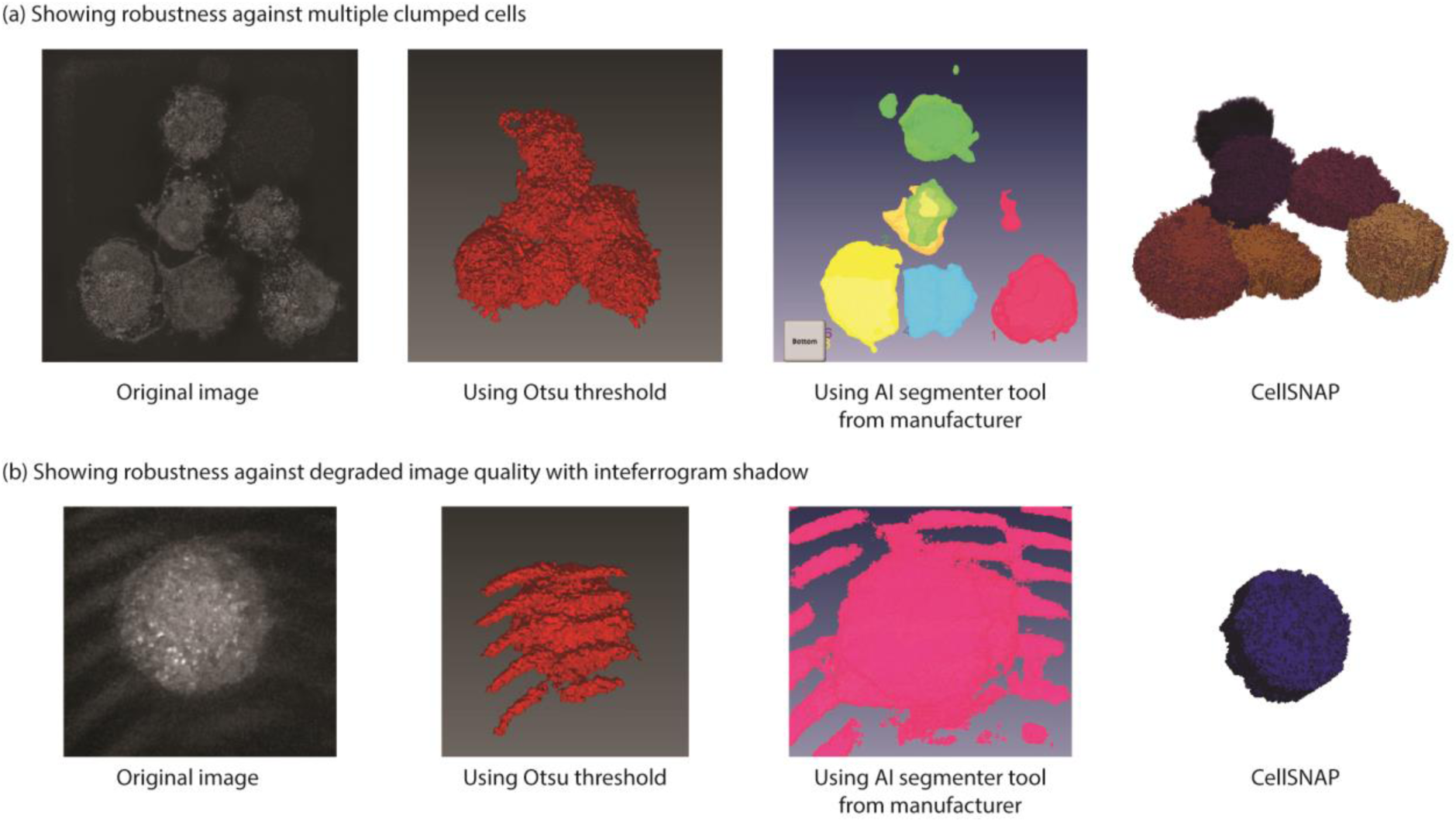
Figure illustrates the efficacy of our algorithm in handling challenging scenarios encountered during QPI imaging. (a) Multiple cells in proximity pose a segmentation challenge for conventional methods such as Otsu threshold and AI-based segmentation. However, our algorithm can accurately segment individual cells even in such clumped conditions. (b) In suboptimal imaging conditions, interferogram noise impedes accurate segmentation by traditional techniques. In contrast, our algorithm is resilient to such noise and can accurately segment cells.

**Table 1:** Computational speed comparison for different methods.

| Method name | Computation time | Total time | System configuration |
| --- | --- | --- | --- |
| Watershed algorithm | File loading – 3 seconds;<br>Processing time ~ 7 seconds | 10 seconds | i7 - 8700K CPU @3.7 GHz – 6 core processor, 64 GB RAM, Nvidia GeForce GTX 1080 - 8GB graphics card |
| AI-enabled segmentation | Running time after training ~ 6 seconds | 6 seconds | i7 - 8700K CPU @3.7 GHz – 6 core processor, 64 GB RAM, Nvidia GeForce GTX 1080 - 8GB graphics card |
| Our algorithm | 2D segmentation ~ 1 second<br>Remaining process ~ 1 second | 2 seconds per cell utilizing a single-core processor. It can be sped up by parallel processors | i5 -7200 @ 2.7 GHz, 16 GB RAM |

In another example, the image captured has interferogram patterns on the image. This typically arises from calibration drift when performing time-lapse over extended periods. Removing this error with just an Otsu threshold proves challenging, given the similarity between pixel values in the interferogram shadows and the cell. A pattern recognition conditional statement may be used to remove this error along with the Otsu threshold, but we are not aware of any implementation of such an algorithm. While AI segmentation tools can be trained with datasets featuring interferogram artifacts, such implementations are computationally intensive and demanding. Moreover, such errors are prevalent for tools based on trained models when presented with unique cases [21]. Introducing enough variations in the training dataset and retraining the algorithm can overcome this, but this process has limitations, as the machine learning algorithm seldom provides details on the methods [22]. Debugging is also difficult if the trained model influences other cases.

The only limitation of our algorithm is that the process assumes that there is only one cell along the z-stack. Therefore, the algorithm would not work for cases where there are multiple cells clumped along the z-axis. Such cases cannot occur for cells cultured in a Petri dish. Multiple cells clumped along the z-axis can occur in the case of spheroids which is currently beyond the scope of our studies.

## Conclusion

QPI is a powerful imaging tool that is making major strides and has found numerous applications in basic biological studies and applied clinical research. As QPI utilizes the optical path length as intrinsic contrast, the imaging is noninvasive and, thereby, allows for monitoring live cell samples over several days without concerns of degraded viability. Therefore, significant recent attention has been focused on developing robust analysis pipelines for quantitative phase images, including the application of convolutional neural networks for computationally substituting chemical stains for cells, extracting biomarkers of interest, and enhancing imaging quality. Yet, 3D segmentation of cells, particularly clumped cells, presents a major challenge, as the existing methods work well only for isolated cells. In this work, we have shown that our cell segmentation algorithm for QPI images outperforms the existing gold standard both in terms of speed and robustness. Our algorithm takes *ca*. 2 seconds per cell on a single-core processor to perform the segmentation. This can easily be parallelized on a multi-core system for further improvement in speed. For the cases where segmentation is possible with the existing standard method, our algorithm has a mean error of 5% for dry mass and 8% for volume measurements. Further morphological analysis, such as the determination of surface area, aspect ratio, circularity, and others, can be done by standard function files on the segmented 3D mask images generated by our algorithm. Therefore, this work can lead to wider adoption of QPI imaging for high-throughput analysis, which was earlier stymied by a lack of suitable cell segmentation tools and lower the barrier to the adoption of QPI imaging modality in biological sciences.

### Cell culture method

U937 cells were seeded in TomoDishes at about 5*10^4^ cells per dish with complete media and PMA (Sigma -Aldrich 79346, 1 uM). The complete media consisted of RPMI-1640 (Gibco, 11875-093), 10% heat-inactivated FBS (Corning, 35-010-CV), 1% P/S, 1%L-Glu (Gibco, 200 mM, 25030081) and 1% HEPES (Gibco, 1M, 15630-080). The seeding day is counted as Day 0.

For M1 polarized cells, on Day 3, PMA media is removed, and cells are washed with PBS. Thereafter, polarizing media consisting of complete media with LPS (List Biological Laboratories, 421, 100 ng/mL) and IFN-y (Sigma-Aldrich, SRP3093, 50 ng/mL) were added to the dish. The cells were incubated for another two days. On day 5, the media was removed, and cells were washed with PBS before fixing.

For M2 polarized cells, on Day 3, PMA media is removed, and cells are washed with PBS. Thereafter, polarizing media consisting of complete media with IL-4 (Sigma-Aldrich, SRP3093, 50 ng/mL) and IL-13 (R&D Systems, 213-ILB-005, 50 ng/mL) were added to the dish. The cells were incubated for another two days. On day 5, the media was removed, and cells were washed with PBS before fixing.

The cells were fixed with 4% PFA at room temperature for 15 minutes. It was then washed with PBS two times. PBS was once again added to fully submerge the fixed cells before imaging.

### Imaging method

The measurements were performed on a Quantitative phase imaging system (HT-1H, Tomocube Inc, Republic of Korea) comprised of 60X water immersion objective (1.2 NA), an off-axis Mach-Zender interferometer with a 532 nm laser, and a digital micromirror device (DMD) for tomographic scanning of each cell [23]. The 3D RI distribution of the cells was reconstructed from the interferograms using the Fourier diffraction theorem as described previously [24]. TomoStudio (Tomocube Inc, Republic of Korea) was used to reconstruct and visualize 3D RI maps and their 2D maximum intensity projections (MIP).

### Code availability

The code for analysis with a sample dataset can be found on the following link: https://github.com/Lconway4C/QPI-cell-segmentation.git

An additional dataset containing RI Tomogram TIFF files can be found on the following link: https://doi.org/10.6084/m9.figshare.23547087

## Acknowledgment

We would like to thank Professor Denis Wirtz and his students Haonan Xu and Bartholomew Starich for their help with macrophage cell culture.

We acknowledge support from the Air Force Office of Scientific Research (FA9550-22-1-0334), National National Institute of General Medical Sciences (1R35GM149272) and the National Cancer Institute (R01-CA238025)

